# Deconvoluting TCR-dependent & -independent activation is vital for reliable Ag-specific CD4^+^ T cell characterization by AIM assay

**DOI:** 10.1101/2024.12.10.627643

**Authors:** Ming Z. M. Zheng, Lauren Burmas, Hyon-Xhi Tan, Mai-Chi Trieu, Hyun Jae Lee, Daniel Rawlinson, Ashraful Haque, Stephen J. Kent, Adam K. Wheatley, Jennifer A. Juno

## Abstract

AIM assays are thought to detect antigen (Ag)-specific T cell responses in an HLA- and cytokine-independent manner. Recent studies using AIM assays identified prominent Th17-like (CCR6^+^) CD4^+^ T cells and circulating follicular T helper cells (cTfh) in anti-viral contexts, but were not observed with peptide/HLA tetramer staining. We demonstrate that CD39^+^ Treg-like and CD26^hi^ Th22-like cells can be activated by cytokines in a TCR-independent manner during *in vitro* Ag stimulation, leading to non-specific upregulation of prototypical AIM readouts. Transcriptional analysis of memory CD4^+^ T cells that underwent TCR-dependent or -independent activation enabled discrimination of *bona fide* Ag-specific T cells from cytokine-activated Treg and Th22 cells. CXCR4 downregulation emerged as a hallmark of clonotypic expansion and TCR-dependent activation in memory CD4^+^ T cells and cTfh. Tracking tetramer-binding cells during re-stimulation showed that CXCR4^-^CD137^+^ cells provide a more accurate measure of the Ag-specific population than standard AIM readouts. This modified assay excludes the predominately CCR6^+^ cytokine-activated T cells that contribute to an average 12-fold overestimation of the Ag-specific population. As AIM assays enable the rapid study of T cell responses against emerging pathogens, our findings provide a highly accurate approach to characterize genuine Ag-specific T cell responses that contribute to protective immunity.

## Introduction

T cells are key effectors of the adaptive immune system that play both direct and indirect roles in protection against pathogens, cancer, and autoimmunity. CD8^+^ T cells have a well described capacity for directed killing of infected or aberrant cells, while CD4^+^ T helper cells coordinate the function of a diverse array of immune mediators, including antigen presenting cells (APCs), CD8^+^ T cells and B cells. Consequently, accurate measurement of the magnitude and quality of antigen (Ag)-specific T cells is critical for a mechanistic understanding of adaptive immunity. Numerous tools have been developed to define the frequency, phenotype, and function of Ag-specific T cells in human clinical samples, each with distinct advantages and caveats. Peptide/MHC multimers ^1^ provide unparalleled precision for the direct identification of Ag-specific T cells but are limited to defined peptide/HLA (pHLA) pairings. Strategies to study a polyclonal Ag-specific T cell response typically rely on *in vitro* re-stimulation by peptides or protein antigens, with responding cells identified using enzyme-linked immunospot (ELISPOT) ^2^, intracellular cytokine staining (ICS) ^3^, and/or proliferation ^4,5^. Recently, surface upregulation of activation-induced markers (AIM) has emerged as a versatile and sensitive approach to study T cell immunity^6–8^, particularly for subsets that produce low levels of cytokines such as follicular helper T cells (and their circulating counterparts, cTfh) ^9^, or regulatory T cells (Treg)^10^.

AIM assays detect upregulation of surface receptors such as OX-40, CD137 (4-1BB), CD25, CD154 (CD40L), PD-L1 (CD274), CD69 or CD200 ^8,9,11–19^ and have been frequently deployed to measure CD4^+^ T cells following infection or vaccination, with widespread use during the COVID-19 pandemic ^14,15,18,20–23^. However, there are inconsistencies in the cellular phenotype of Ag-specific cells identified by AIM versus those defined using pHLA tetramers or ICS. For example, we and others have reported that CD4^+^ T cells responding to SARS-CoV-2 spike, HIV gag, hepatitis B surface Ag and tetanus in the AIM assay include a substantial population of CCR6^+^ Th17-like CD4^+^ T cells ^14,20,24–26^. Nevertheless, IL-17 production is rarely detected in response to stimulation with these Ags ^14,20^, and cells bound by pHLA tetramers are almost exclusively CCR6^-^ ^16,27^. Such inconsistencies confound the ability to precisely quantify T cell responses and define qualitative features associated with protective function.

We therefore sought to characterize the precise relationship between the Ag-specific CD4^+^ T cells defined using AIM assays versus pHLA tetramers. Using a combination of transwell experiments and transcriptional analysis, we identified two populations of memory T cells, CD39^+^ Treg-like and CD26^hi^ Th22-like cells, that are enriched for CCR6 expression and upregulate common AIM markers in a TCR-independent fashion *in vitro*. These populations could be differentiated from *bona fide* Ag-specific T cells based on gene expression, the presence of clonal expansion, and tetramer binding. Using differential gene expression analysis, we screened alternative AIM readouts and determined that identifying responsive cells via CXCR4 downregulation and CD137 upregulation improved the precision of the AIM assay. Our study provides further refinement and standardization of assays to measure T cell immunity, which is critical for informative studies in clinical immunology to support assessment of vaccines and treatments for disease.

## Results

### AIM assays and pHLA class II tetramers identify discordant populations of Ag-specific CD4^+^ T cells

Prominent populations of CCR6^+^ SARS-CoV-2 spike (S)-specific CD4^+^ T cells have been reported by multiple groups using OX-40/CD25/CD137-based AIM assays ^14,20,24,25^. In contrast, CCR6 surface expression is largely lacking on S-specific cells identified using pHLA tetramers ^16,27^. Here, we directly compared the phenotype of CD4^+^ T cells recognizing spike S_751-766_ or S_167-180_ epitopes identified by AIM or tetramer. In line with past reports, we find AIM^+^ cells (defined as either OX-40^+^CD25^+^ or OX-40^+^CD137^+^) were predominately CCR6^+^, while tetramer-binding cells were almost exclusively CCR6^-^ (**Fig. 1A-B**). We noted a similar phenomenon in individuals receiving seasonal influenza vaccination, where HA-specific CD4^+^ T cells identified by AIM included a prominent CCR6^+^ population (**Fig. S1A-B**), in contrast to the published CCR6^-^ phenotype of cells binding HA tetramers ^28,29^.

**Figure 1.**
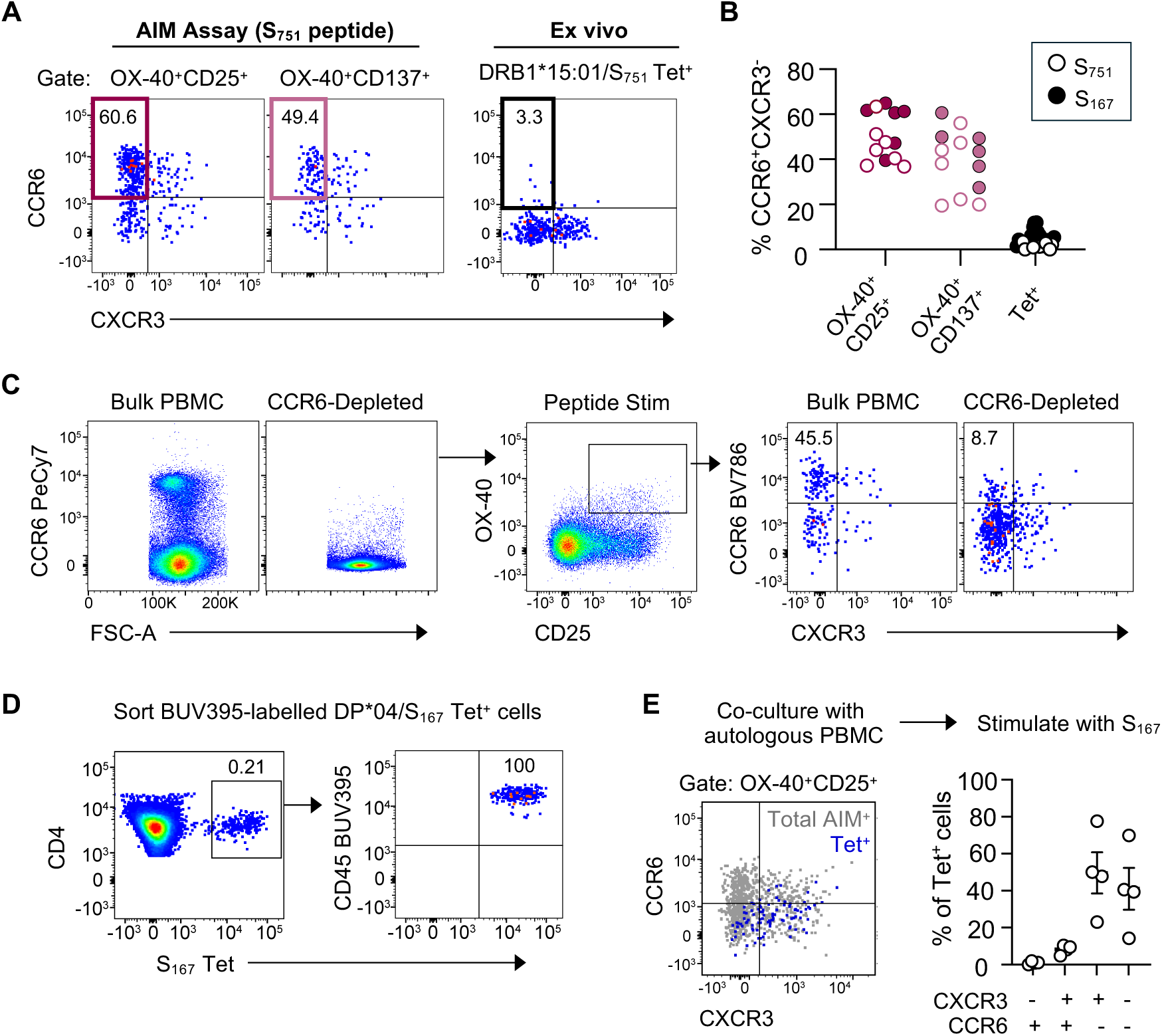
CCR6^+^ spike-specific CD4^+^ T cells are not detected by pHLA tetramers. **(A)** Representative staining of CCR6 and CXCR3 expression on S_751-766_–specific memory CD4^+^ T cells in a single donor as determined by AIM assay after *in vitro* peptide stimulation or identified by DRB1*15:01/S_751_ tetramer *ex vivo.* **(B)** Frequencies of CCR6^+^CXCR3^-^ S_751_- or S_167_-specific memory CD4^+^ T cells defined by AIM or tetramers. For the S_751_ epitope, data from n=6 individuals and representative of the respective cohorts (n=20-40) derived from Juno *et al.* ^20^ and Wragg *et al.* ^27^. For the S_167_ epitope, data from n=7 individuals. **(C)** Expression of CCR6 on AIM^+^ memory CD4^+^ T cells following *in vitro* S2 peptide pool stimulation of untouched or CCR6-depleted PBMCs from SARS-CoV-2 vaccinated subjects. Data representative of two independent experiments. **(D)** Verification of CD45 BUV395 labelling of sorted DPB1/04*01-S_167_ tetramer^+^ CD4^+^ Tmem. **(E)** Representative expression of CCR6 and CXCR3 on total AIM^+^ Tmem (grey) or CD45 BUV395^+^ Tmem (blue) following *in vitro* stimulation with S_167_ peptide. Graph indicates the proportion of CD45 BUV395^+^ cells with each phenotype, from n=4 individuals.

To investigate whether CCR6 expression was impacted by *in vitro* peptide stimulation, we cultured PBMCs from SARS-CoV-2-vaccinated individuals with a spike peptide pool. In samples subjected to prior depletion of CCR6^+^ cells, we observed minimal *de novo* CCR6 surface expression on AIM^+^ cells (**Fig. 1C**). We next sorted HLA-DPB1*04:01/S_167_ tetramer-specific T cells, labelled them with CD45-BUV395 antibody, and cultured them in the presence of cognate peptide and autologous PBMCs (**Fig. 1D**). After 20 hrs of stimulation, tetramer^+^ T cells remained almost exclusively CCR6^-^ (**Fig. 1E**), confirming that CCR6 upregulation is not artifactually induced via the stimulation conditions of the AIM assay.

### CCR6^+^ AIM^+^ cells lack clonal expansion following Ag re-exposure

To further explore the heterogeneity of AIM^+^ cells, we transcriptionally profiled CD4^+^ T cells in three individuals receiving seasonal influenza vaccination. PBMCs collected at baseline, day 7, and day 21 post-immunization were stimulated *in vitro* with recombinant HA proteins. Memory CD4^+^ T cells responsive (OX-40^+^) or non-responsive (OX-40^-^) to stimulation were sorted for transcriptional analysis using the 10x Genomics Chromium scRNAseq platform. Following dimensionality reduction by UMAP, activated cells were identified based on surface protein expression of CD25 and/or OX40 via antibody-derived tags (ADTs) (**Fig. S1C**). Profiling of the AIM^+^ cells identified 5 putative clusters based on differential gene expression (**Fig. S1D**). Interestingly, TCR sequencing indicated that clonally expanded T cell populations were present only within cluster 4, which showed minimal expression of CCR6 RNA or protein (**Fig. S1E**). These cells also exhibited relatively low expression of *CXCR4* (**Fig. S1E**), a feature associated with TCR-mediated activation ^30,31^. Cluster 3 exhibited classical Treg gene signatures with enrichment of *FOXP3, IL2RA, CTLA4*, *IKZF2* and *IKZF4* (**Fig. S1E-F**). Gene expression patterns for clusters 0, 1 and 2 were less cell-type distinctive (**Fig. S1F**), without strong overlap with easily defined CD4^+^ T cell subsets. Overall, our data suggest that in addition to clonally expanded Ag-specific CD4^+^ T cells, the AIM assay identifies a mixed population of Treg and other CD4^+^ T cell subsets that may become activated in an Ag-independent manner.

### A subset of CCR6^+^ CD4^+^ T memory cells are prone to IL-2 mediated activation

To address whether common AIM markers (OX-40, CD25, CD137, CD154) are upregulated in the absence of TCR-mediated stimulation or cell-cell contact, we adapted an *in vitro* stimulation and transwell culture system ^8,12,32^. PBMCs seeded in the bottom chamber were stimulated via anti-CD3/CD28 Dynabeads, while autologous PBMCs served as ‘sentinel’ cells in the upper chamber (**Fig. 2A**). Sentinel CD4^+^ T cells upregulated all combinations of AIM markers upon exposure to the Dynabead chamber relative to unstimulated controls **(Fig. 2B-C**). Activated sentinel cells were almost exclusively CD45RA^-^ memory (**Fig. 2D**), comprised both cTfh (CXCR5^+^) and non-cTfh subsets (**Fig. 2D**), and were substantially enriched for CCR6 expression (**Fig. 2E**).

**Figure 2.**
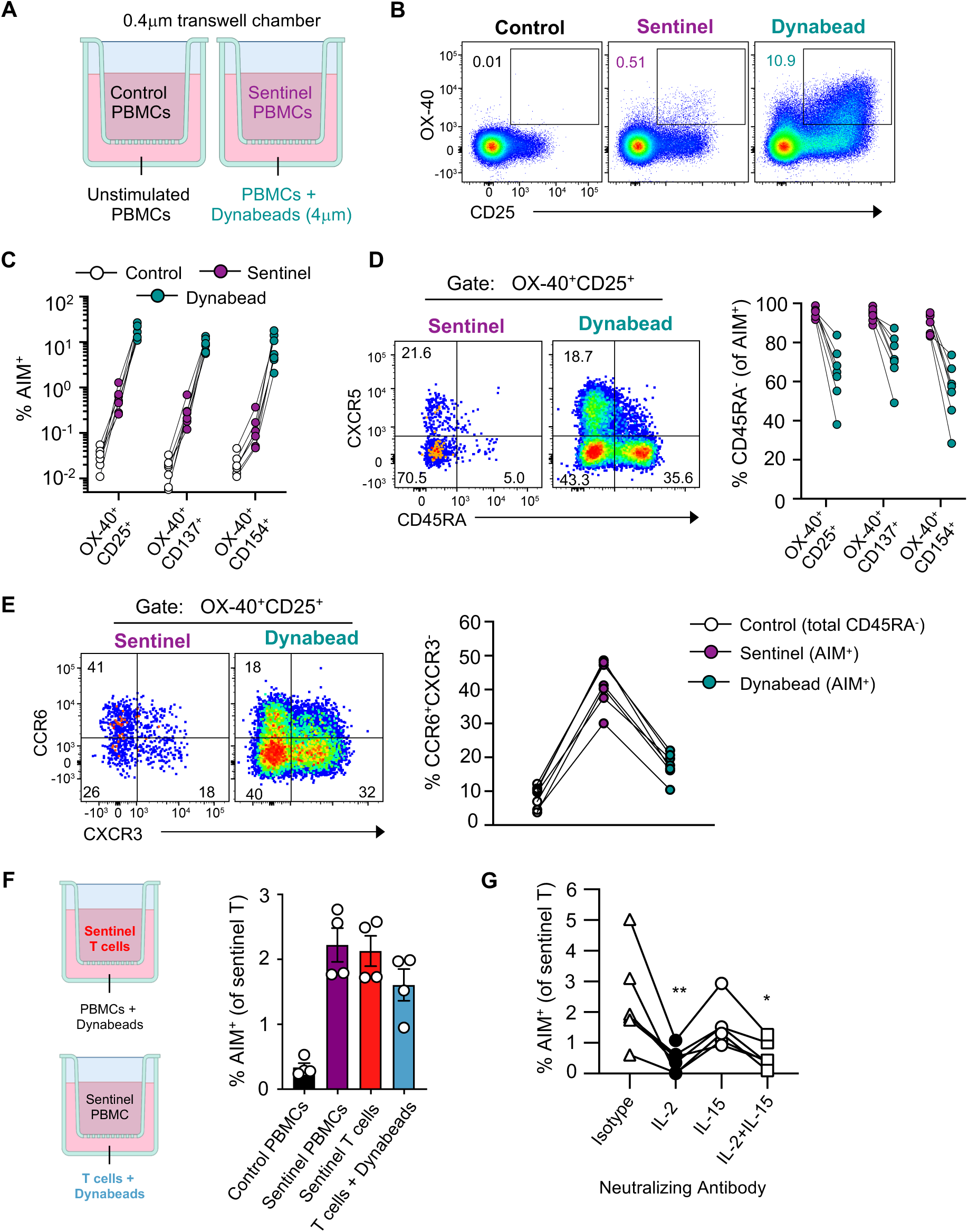
Soluble factors acting directly on T cells contribute to TCR-independent activation of CCR6^+^ CD4^+^ Tmem. **(A)** Schematic of transwell stimulation assay. **(B)** FACS profiles and **(C)** frequency of AIM^+^ CD4^+^ Tmem in control, sentinel (purple) or Dynabead (teal) chambers, from n=7 individuals. **(D)** Distribution of naïve (CD45RA^+^), memory (CD45RA^-^ CXCR5^-^) and cTfh (CD45RA^-^CXCR5^+^) cells among AIM^+^ cells in the sentinel and Dynabead wells with corresponding frequencies (n=7 individuals). **(E)** Frequency of AIM^+^ events with a CCR6^+^CXCR3^-^ phenotype among the total CD45RA^-^ population of the control chamber, or AIM^+^CD45RA^-^ population of the sentinel and Dynabead chambers (n=7 individuals). **(F)** AIM^+^ frequencies on sentinel CD4^+^CD45RA^-^ populations of PBMCs, enriched T cells, or a sentinel population of PBMCs that were exposed to Dynabead-stimulated T cells (n=4 individuals). **(G)** Proportion of AIM^+^ cells in the sentinel well (enriched T cell condition) following incubation with isotype control, anti-IL-2, anti-IL-15, or anti-IL-2/IL-15 antibodies in the lower chamber (n=5 individuals). Statistics assessed by Friedman test with Dunn’s multiple comparison test comparing against the isotype group. \**p* < 0.5, \*\**p* < 0.01.

To determine whether soluble factors were acting directly on the sentinel T cells or via an intermediate cell type, we substituted enriched T cells as either the sentinel or Dynabead-stimulated population (**Fig. 2F**). Activated sentinels were observed irrespective of whether the upper chamber contained complete PBMCs or isolated T cells (**Fig. 2F**). Similarly, Dynabead-stimulated T cells in the bottom chamber were sufficient to drive sentinel activation, albeit at levels slightly reduced (1.3-fold) compared to total PBMCs (**Fig. 2F**). Thus, secretion of a soluble factor produced by T cells alone is sufficient to drive upregulation of OX-40, CD25, CD137 and CD154 on closely localized cells. Consistent with prior reports of OX-40 and CD154 upregulation on memory CD4^+^ T cells by γ−chain cytokines ^33–35^, neutralization of either IL-2 and IL-15 in the lower chamber of the transwell system decreased sentinel T cell activation by 5.4- or 1.5-fold respectively, albeit with no synergistic effect (**Fig. 2G**). Thus, IL-2 acting directly on T cells appears to be a primary driver of non-specific AIM upregulation *in vitro*.

### Transcriptional profiling identifies TCR-independent activation of Treg and CD26^hi^ Th22-like cells

We next used bulk RNAseq to clarify the identity of the CCR6^+^ IL-2-responsive T cells in the transwell system. CXCR3^+^, CCR6^+^ or CXCR3^-^CCR6^-^ subpopulations of Dynabead-activated CD4^+^ T cells were sorted based on OX40/CD25 co-expression as reference controls for TCR-based stimulation (**Fig. S2A**). From the sentinel population, non-activated CXCR3^+^, CCR6^+^ or CXCR3^-^CCR6^-^ subpopulations were sorted as cells refractory to cytokine-dependent activation to serve as comparators for the activated sentinel CD4^+^ T cells (CCR6^+^ OX40^+^CD25^+^ cells) (**Fig. S2A**).

Consistent with our flow cytometry data, both Dynabead and sentinel activated cells expressed high levels of the common AIM readouts *IL2RA* (CD25), *TNFRSF4* (OX40), *TNFRSF9* (CD137), *CD40LG* (CD154), *CD274* (PD-L1) and *CD69* (CD69) ^9,19^ (**Fig. 3A**, **Fig. S2B**). Relative to controls, CCR6^+^AIM^+^ sentinel cells were enriched for Treg-associated genes (*FOXP3*, *TNFRSF18*, *CD74*) (**Fig. 3A**, **Fig. S2C**), as well as *DPP4* (CD26), *IL22* (IL-22) and *IL7F* (IL-17F) (**Fig. 3A**, **Fig. S2D**). The latter set of genes are characteristic of Th17/Th22 cells, which have been described as a cytokine-responsive, IL-17/IL-22 secreting population of CCR6^+^ CD26^hi^ memory CD4^+^ T cells in humans ^36^. We validated these phenotypes at the protein level using flow cytometry, where we observed high frequencies of both CD39^+^ Tregs ^37,38^ and CD26^hi^ cells within sentinel AIM^+^ T cells (**Fig. 3B-C**). Together, these populations accounted for ∼78% (IQR 75.43 to 82.13) of cytokine-responsive sentinel cells.

**Figure 3.**
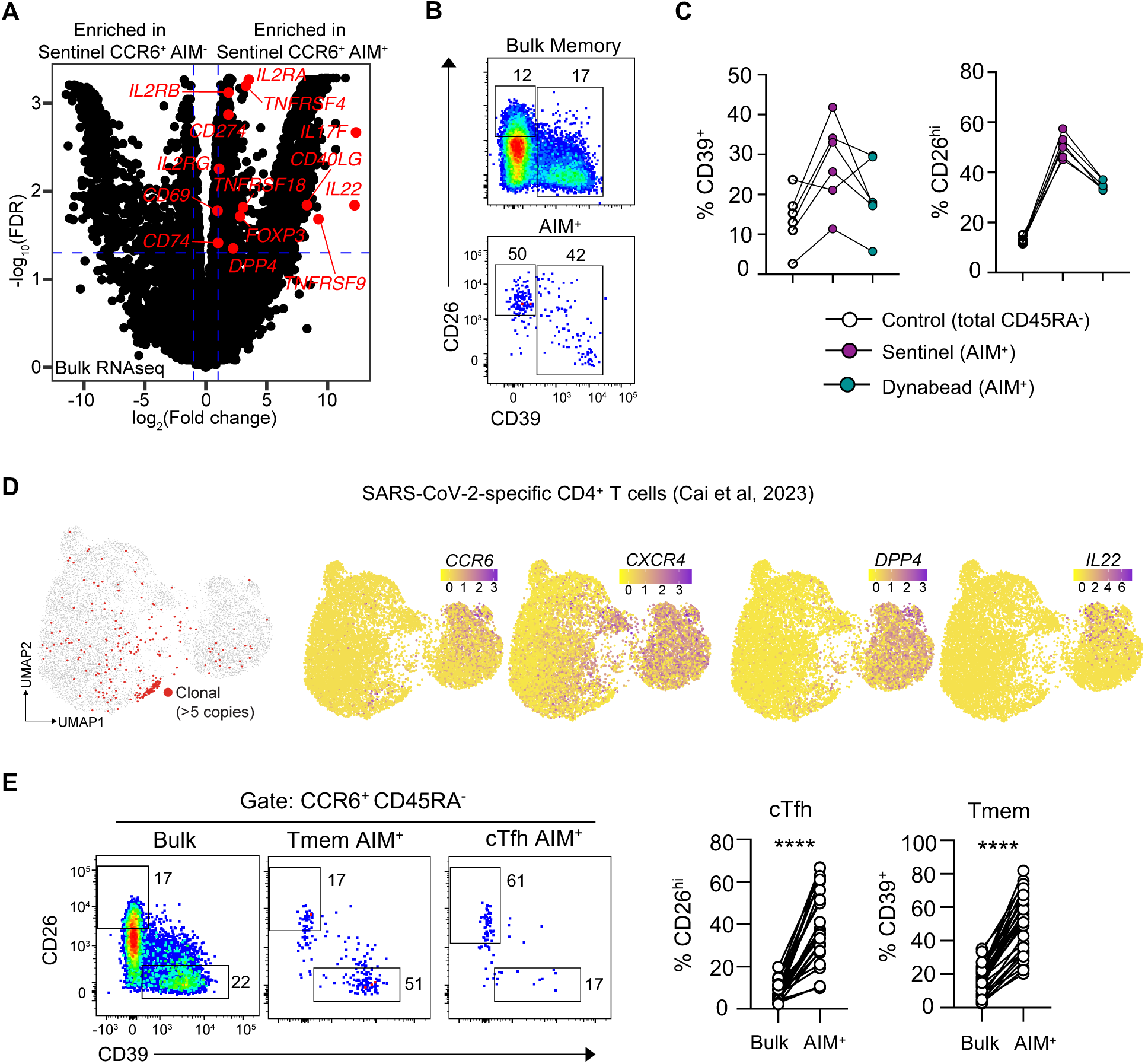
CD26^hi^ Th22-like and CD39^+^ Treg-like cells are enriched within CCR6^+^ AIM^+^ CD4^+^ T cells. **(A)** Volcano plot of selected genes enriched in sentinel AIM^+^ CCR6^+^ cells relative to sentinel AIM^-^ CCR6^+^ CD4^+^ T cells identified by bulk RNAseq (n=1 individual). Dotted line indicates criteria threshold of -log_10_(0.05) FDR and at least a log_2_ fold change of - 1 or 1. **(B)** FACS plots and **(C)** frequency of CD26^hi^CD39^-^ and CD39^+^ populations among bulk or AIM^+^ (CD25^+^OX40^+^) Tmem (n=6 individuals). **(D)** UMAP plots of the single cell sequencing dataset from (Cai *et al.*, 2023) ^24^ showing positioning of expanded TCR CDR3 clonotypes shared between S-specific AIM^+^ memory CD4^+^ T cells, or expression of *CCR6* (CCR6), *DPP4* (CD26), *IL22* (IL-22) and *CXCR4* (CXCR4). **(E)** AIM responses among PBMCs collected from a randomized clinical trial of mRNA-1273.214 WT/BA.1 COVID-19 bivalent booster vaccination (n=33 individuals). FACS plots and graphs show frequencies of CD26^hi^CD39^-^ and CD26^-^CD39^+^ cells among bulk or AIM^+^ subsets of CCR6^+^ cTfh and Tmem populations. Statistics assessed by two-tailed Wilcoxon matched-pairs signed rank test. \*\*\*\**p*<0.0001.

To determine whether cytokine-driven activation of Treg and CD26^hi^ cells was the cause of the discrepancy between AIM and tetramer data (**Fig. 1**), we sought validation of these signatures in the context of SARS-CoV-2 spike peptide-based AIM assays. We first examined a publicly available transcriptional dataset of S-specific AIM^+^ CD4^+^ T cells from convalescent donors ^24^, which exhibited two major clusters of activated cells (**Fig. 3D**). Consistent with our transcriptomic profiling of HA-specific AIM^+^ cells (**Fig. S1**), we found clonally expanded S-specific T cells did not express *CCR6* and exhibited low expression of *CXCR4* (**Fig. 3D**). A separate population of cells exhibited the cytokine-driven gene expression signature, including *CCR6*, *DPP4*, and *IL22* (**Fig. 3D**). Secondly, we stimulated PBMCs from a COVID-19 mRNA booster vaccine trial ^39^ with a pool of spike peptides. Both CD26^hi^ and CD39^+^ Treg were significantly enriched within AIM^+^ CCR6^+^ cells (**Fig. 3E**). The enrichment of CD26^hi^ cells was particularly prominent within the cTfh compartment (4.7-fold compared to total CCR6^+^ cTfh), while Treg were highly enriched among CXCR5^-^ memory T cells (3.2-fold vs total CCR6^+^ Tmem) (**Fig. 3E)**. Collectively, these data indicate that cytokine responsive Tregs and Th17/Th22 cells are a major contributor to populations designated as ‘Ag-specific’ T cells identified using AIM assays.

### Differential gene expression analysis identifies specific markers of TCR-dependent activation

Although the cytokine-responsive cells are highly enriched for CCR6 expression, they are not exclusively CCR6^+^ (**Fig. 2E**), precluding the possibility of simply gating out CCR6^+^ cells when analyzing AIM datasets. We therefore sought to identify activation markers that are more reliably indicative of TCR-dependent activation. We parsed the transwell RNAseq dataset for candidate genes with known immune function and surface expression amenable to detection by flow cytometry, and identified *CXCR4* (downregulated by Dynabead activation), *LY75* (CD205/DEC-205) and *TNFRSF8* (CD30) (upregulated by Dynabead stimulation) (**Fig. S3A-B**). We screened permutations of these and existing AIM markers in the transwell assay to identify combinations that detected Dynabead-mediated activation but showed minimal perturbation among the sentinel cTfh and Tmem populations (see Materials & Methods) (**Fig. S3C-D**).

We next determined whether these combinations were suitable to identify Ag-specific T cells following 15mer peptide stimulation. PBMCs from SARS-CoV-2 convalescent/vaccinated individuals were stimulated with a spike peptide pool or SEB as a positive control. Screening revealed that 11 out of the 19 new AIM combinations were insufficiently sensitive to detect any S-specific CD4^+^ Tmem or cTfh responses (**Fig. S3E-F**). Of the remaining permutations, the combination of CXCR4 downregulation and OX-40 or CD137 upregulation most reliably identified a population of S-specific CD4^+^ Tmem and cTfh with reduced CCR6^+^CXCR3^-^ frequencies (**Fig. 4A**) compared to the standard AIM. A direct comparison of the cells identified by conventional (CD25^+^OX40^+^) or modified AIM (CXCR4^-^CD137^+^) revealed that a similar frequency of activated CXCR3^+^ cTfh or Tmem were detected by both approaches (**Fig. 4B-C**). In contrast, the total frequency of activated CCR6^+^ cells dropped 19.8- and 48.9-fold (for cTfh and Tmem, respectively) when using the modified AIM gate (**Fig. 4B-C**). Importantly, this does not reflect an intrinsic inability of CCR6^+^ cells to acquire the CXCR4^-^ CD137^+^ phenotype after stimulation, as SEB comparably activated all CCR6/CXCR3 T cell subsets (**Fig. 4A**).

**Figure 4.**
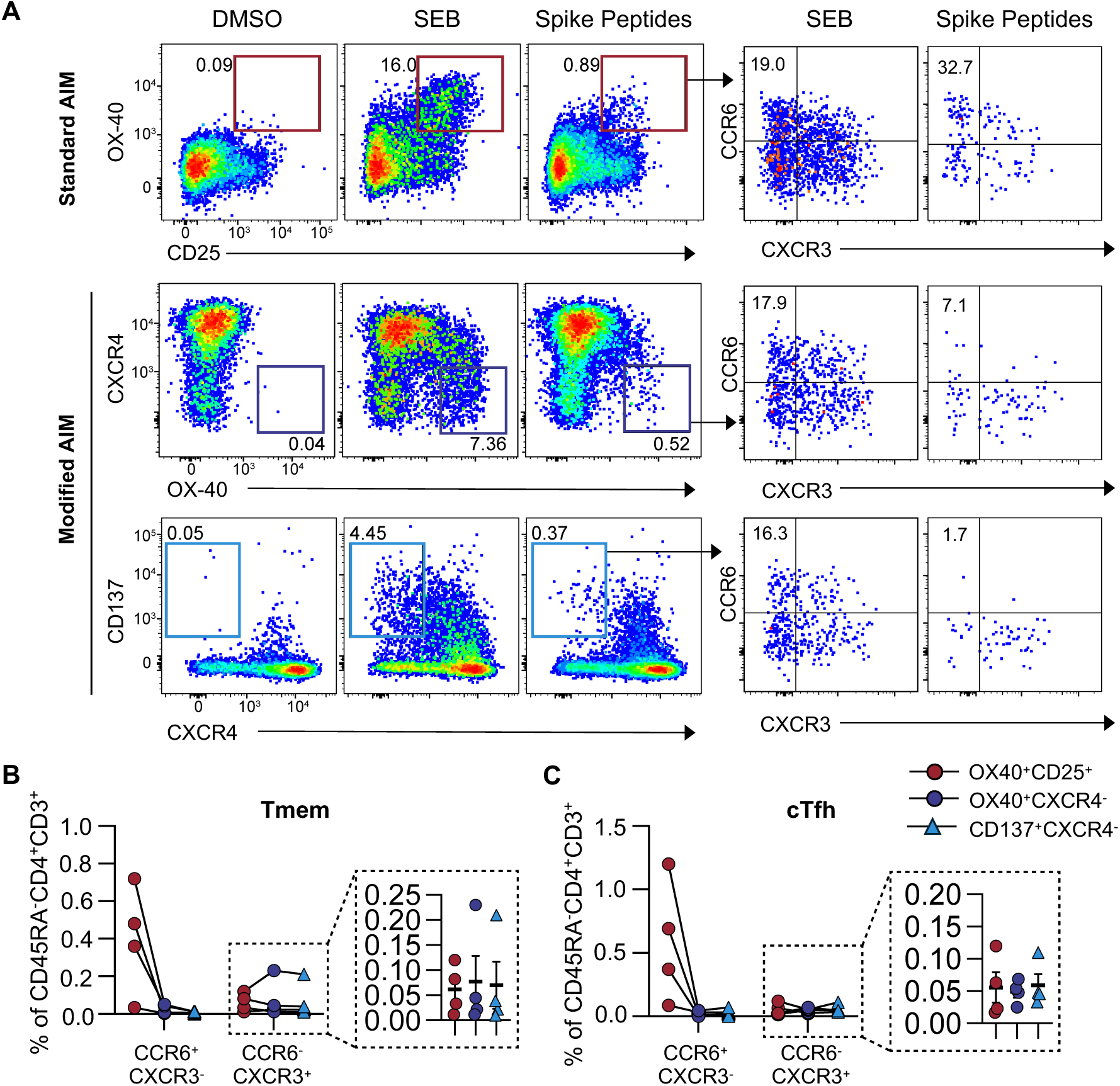
Measurement of CXCR4 downregulation during *in vitro* stimulation provides an alternate AIM readout. PBMCs from SARS-CoV-2 vaccinated/infected individuals were stimulated with a peptide pool spanning the S2 region of the spike protein or the positive control SEB. **(A)** Representative staining of standard (OX-40^+^CD25^+^) or modified (CXCR4^-^OX-40^+^ or CXCR4^-^CD137^+^) AIM responses in a single individual. Peptide-stimulated AIM^+^ cells were further assessed for CCR6 and CXCR3 co-expression. **(B-C)** Abundance of AIM^+^ CCR6^+^CXCR3^-^ and CCR6^-^CXCR3^+^ cells expressed as a percentage of total **(B)** Tmem or **(C)** cTfh subsets post peptide stimulation (n=4 individuals).

### CXCR4^-^CD137^+^ AIM readout demarks Ag-specific CD4^+^ Tmem identified by pHLA tetramers

To validate that the modified AIM reliably identifies Ag-specific CD4^+^ T cells, we isolated and BUV395-labelled S_167_ tetramer^+^ T cells from COVID-19 vaccinated individuals, then re-stimulated them in culture with autologous PBMCs as in **Fig. 1E**. Following S_167_ peptide stimulation, both conventional and modified AIM gates identified similar numbers of tetramer^+^ cells, indicating comparable sensitivity (**Fig. 5A-B**). However, tetramer^+^ cells constituted only 6% of the total population of CD25^+^OX40^+^ cells within the culture (**Fig. 5C**), highlighting high levels of cytokine-driven activation that resulted in an over-representation of CCR6^+^ and CXCR3^-^CCR6^-^ cells. In contrast, the modified AIM exhibited substantially higher specificity, with tetramer^+^ cells comprising 31% (CXCR4^-^OX40^+^) to 43% (CXCR4^-^CD137^+^) of the new AIM gate (**Fig. 5C**). As a result, the abundance of Ag-specific CD4^+^ Tmem measured by new (CXCR4^-^CD137^+^ or CXCR4^-^OX40^+^) but not conventional (CD25^+^OX40^+^) AIM readouts were more comparable to those identified by tetramer in both frequency (**Fig. 5D**) and CCR6/CXCR3 phenotype (**Fig. 5E**). Thus, a modified AIM assay that defines activated CD4^+^ T cells as CXCR4^-^CD137^+^ increases the specificity of the assay without compromising sensitivity.

**Figure 5.**
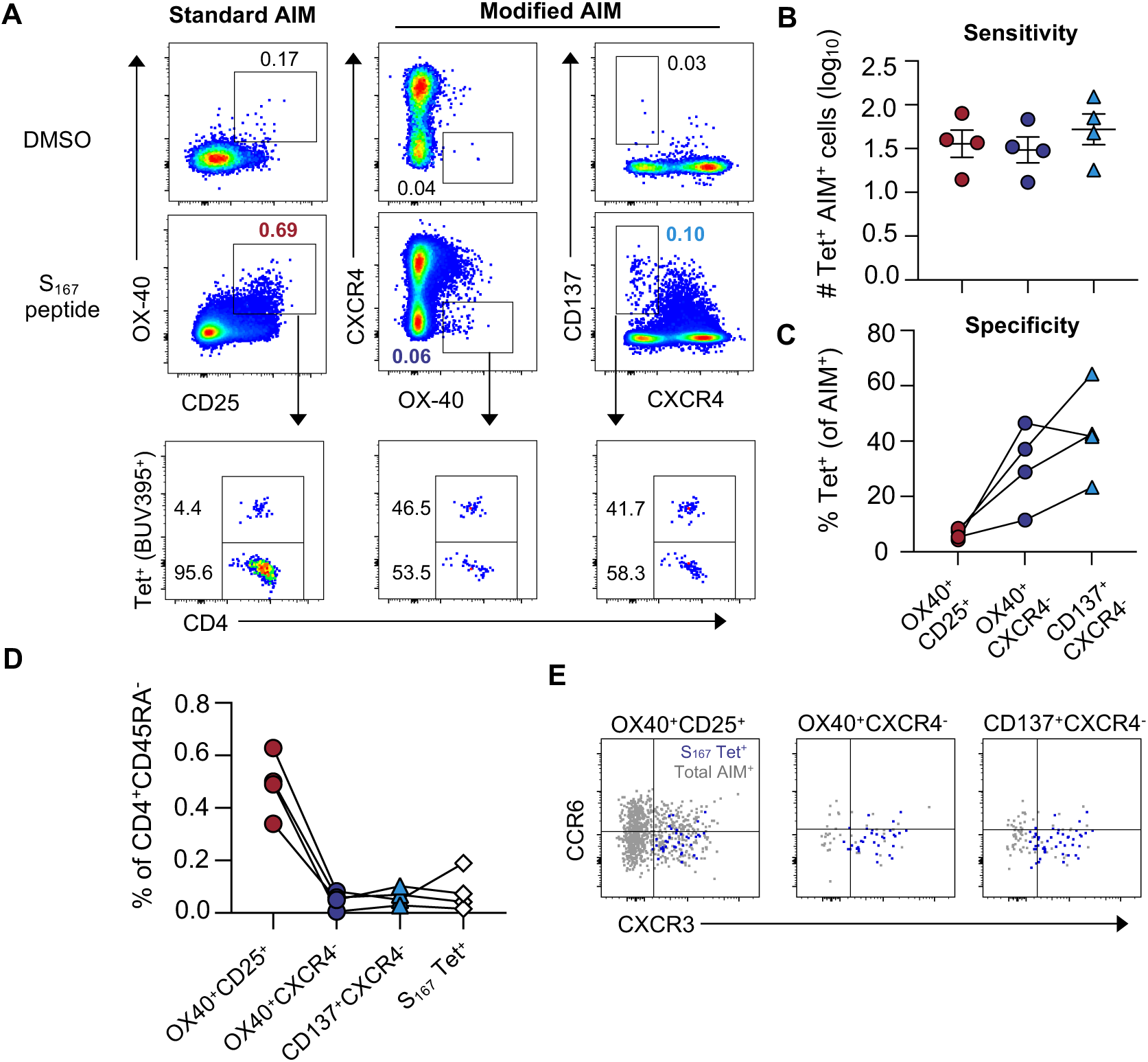
The use of CXCR4^-^CD137^+^ as an AIM readout enables the specific identification of Ag-specific CD4^+^ Tmem. CD45 BUV395-labelled DPB1/04*01-S_167_ tetramer^+^ Tmem were added back into unlabeled autologous PBMCs and stimulated *in vitro* with S_167_ peptide. **(A)** Representative staining of AIM^+^ cells using standard (OX40^+^CD25^+^) and modified (CXCR4^-^ OX40^+^ and CXCR4^-^CD137^+^) AIM gates. The proportion of BUV395 labelled cells within each AIM^+^ population is shown. **(B)** Absolute numbers of tetramer^+^ (BUV395^+^) cells identified by each AIM readout. **(C)** Frequency of tetramer^+^ cells within each AIM gate. **(D)** Frequencies of S_167_-specific CD4^+^ Tmem as defined by AIM combination or pHLA tetramer. n=4 individuals for **(B)-(D). (E)** Representative expression of CCR6 and CXCR3 on total AIM^+^ (grey) and AIM^+^ tetramer^+^ (blue) populations (representative of 4 individuals).

## Discussion

Accurate identification of Ag-specific CD4^+^ T cells is critical to understanding features of protective immunity ^40^. We provide clear evidence that conventional AIM identifies a combination of Ag-specific and cytokine-responsive CD4^+^ T cells. Importantly, we find that cytokine-dependent activation is largely restricted to two populations of memory CD4^+^ T cells: Tregs and CD26^hi^ Th17/Th22-like cells. Accordingly, we and others find no evidence for non-specific activation of naïve CD4^+^ T cells ^8,9,12^, or of unrelated epitope-specific CD4^+^ T cells identified by pHLA tetramers ^9^. The comparatively high expression of CCR6 on CD26^hi^ cells and some Treg explains the appearance of Th17-like cells that have been reported in many AIM studies, but not detected by ICS or tetramer. Overall, our data suggest that when Ag-specific T cells are activated *in vitro* with protein or peptides, they secrete cytokines (including IL-2) that drive TCR-independent activation of Treg and Th22 cells. In the absence of Ag-specific T cells, no cytokine signals are produced, explaining the overall specificity of the assay and its ability to discriminate between Ag-exposed and -unexposed individuals ^9,20,41^.

The existence of cytokine-responsive CD4^+^ Tmem has been well described; combinations of IL-2, IL-7, IL-15, TNF, IL-6 and IL-10 have all been demonstrated to drive proliferation and/or CD69, OX-40, and CD154 expression on human CD45RO^+^ T cells *in vitro* ^33–35^. Further characterization of specific memory subsets targeted by each cytokine is generally lacking, but our data suggest a dominant role for IL-2 in the activation of both Treg and Th22-like CD4^+^ Tmem. It is known that the prototypical AIM combination of CD25^+^OX40^+^ detects activation of CD39^+^ Tregs ^10,42^, many of which express CCR6 at steady state ^43^ or upregulate CCR6 upon stimulation ^44^. Cytokine-driven activation of these cells is consistent with observations that IL-2 can induce FoxP3 expression in CD4^+^ T cells independently of direct TCR stimulation ^45^ and that human Tregs can respond to exogenous IL-2 *in vitro* ^43^. In addition to Tregs, we found substantial cytokine-dependent activation of CCR6^+^CD26^hi^ cells, which have previously been shown to produce IL-17 and IL-22 ^36,46–48^. Co-expression of CCR6 and CD26 is a hallmark of innate-like T cells, including cytokine-responsive CD161^+^ CD8^+^ T cells, MAIT cells and Vδ2^+^ T cells ^49,50^. Further work will be required to elucidate the precise functions and mechanisms of activation for CD4^+^ CD26^hi^ Th17/Th22 cells.

An ideal AIM assay would identify only cells activated through TCR-dependent signalling, without the *a priori* exclusion of, for example, all Th17 cells. By running an extensive screen of genes that distinguished TCR-dependent and -independent activation, we identified CXCR4^-^CD137^+^ as a reliable AIM combination that maintained sensitivity but improved specificity. The downregulation of CXCR4 is a feature of T cell activation ^30,31,51^, resulting in decreased responsiveness to CXCL12 and redirection of cells to inflamed tissues rather than lymph nodes or the bone marrow ^52^. Conversely, cytokines (including IL-2) are known to upregulate CXCR4 on T cells *in vitro* ^31,53,54^. Together, these observations imply that CXCR4 downregulation provides a tractable path forward to facilitate accurate identification of Ag-specific T cells in diverse contexts, including clarifying the CCR6^+^ Th17-like Tmem reported in malaria or *Mycobacterium tuberculosis* infections ^55–57^.

The implications of TCR-independent activation in the AIM assay are three-fold. First, the misidentification of cytokine-activated T cells overestimates (by an average of 12-fold) the magnitude of the Ag-specific T cell response. As the AIM assay is regularly used for assessing vaccine responses ^13,16,19,58^, this overestimation will impact the underlying cellular immunogenicity of a given vaccine. Second, a cytokine-activated CD4^+^ Tmem population will confound RNAseq experiments seeking to define expression profiles of Ag-specific T cells, although single cell sequencing approaches provides a pathway to exclude such cells. Third, the prominent phenotype of cytokine-activated T cells confounds studies to identify CD4^+^ T cell subsets that best correlate with protective immune responses. We and others have inferred differences in cTfh quality (i.e. correlation to neutralizing Abs) and durability based on the skewing of AIM^+^ cTfh toward or away from a CCR6^+^ phenotype ^14,20,25^. Instead, the relative proportion of AIM^+^ cells that are CCR6^+^ more likely depends on the magnitude of the *bona fide* Ag-specific T cell response; when Ag-specific T cell frequency is low, the majority of AIM^+^ cells are likely to be cytokine-responsive and therefore CCR6^+^ biased, explaining why the frequency of cTfh17 cells is reported to increase over time ^14,25^. These observations underscore the need to accurately identify Ag-specific T cell responses to derive insights that inform optimal vaccine-mediated protection.

In summary, our findings demonstrate classical AIM assay readouts (including OX-40, CD137 and CD154) erroneously include substantial populations of cytokine-activated CCR6^+^ CD39^+^ Treg and CD26^hi^ Th17/Th22-like CD4^+^ Tmem. The inclusion of CXCR4 downregulation as an alternative AIM combination can more faithfully recapitulate the frequency and phenotype of Ag-specific T cells identified by pHLA tetramers, allowing more accurate study of cellular immunity responses against pathogens.

## Methods

### Participant recruitment and sample collection

The study protocols were approved by the University of Melbourne Human Research Ethics Committee (13793, 23497 and 2024-21198-48733-4) and Royal Melbourne Hospital Ethics Committee (study number 2021/272), and all associated procedures were carried out in accordance with the approved guidelines. All participants provided written informed consent in accordance with the Declaration of Helsinki. Patients who had recovered from COVID-19 and/or been vaccinated with BNT162b2 or mRNA-1273 were recruited through contacts with the investigators and invited to provide a blood sample. Samples from individuals receiving the mRNA-1273.214 bivalent vaccine were collected as part of a randomized, open-label clinical trial (ACTRN12622000411741) ^39^. No statistical methods were used to predetermine sample sizes. For all participants, whole blood was collected with sodium heparin anticoagulant. PBMCs were isolated via Ficoll-Paque (Sigma Aldrich), cryopreserved in 90% fetal calf serum (FCS)/10% dimethyl sulfoxide (DMSO), and stored in liquid nitrogen.

### Generation of peptide MHC II tetramers

Human DPB1*04:01 SSANNCTFEYVSQPFLMDLE (SARS-CoV-2 S_167_) biotinylated monomers were generated by NIH Tetramer Core Facility. DRB1*15:01 NLLLQYGSFCTQLNRAL (SARS-CoV-2 S_751_) biotinylated monomers were produced by ProImmune. Biotinylated monomers were tetramerized by sequential addition of streptavidin-PE (Thermo Fisher).

### Tetramer staining

Cryopreserved PBMCs were thawed in RPMI-1640 (Thermo Fisher) with 10% FCS and 2% penicillin-streptomycin (RF10), washed and counted. Up to 10e6 PBMCs were washed in 2% FCS/PBS before incubation in 50 nM dasatinib (in 2% FCS/PBS) at 37°C for 30 mins. Tetramers were then added at 4 µg/ml at 37°C for 60 mins. Cells were washed in PBS, stained with Live/Dead Fixable Aqua stain (Thermo Fisher), and incubated for 30 mins at 4°C with a surface stain antibody cocktail containing CD20 BV510 (2H7, BD Biosciences), CD3 BUV805 (SK7, BD Biosciences), CD4 BV605 (RPA-T4, BioLegend), CD45RA R718 (HI100, BD Biosciences), CXCR5 PE-Cy7 (MU5UBEE, Thermo Fisher), CXCR3 PE-Dazzle594 (G025H7, BioLegend), CCR6 BV785 (G034E3, BioLegend). Cells were then washed with 2% FCS/PBS and fixed with Cytofix (BD Biosciences), with data acquisition on an LSR Fortessa (BD Biosciences). Data were analyzed using FlowJo (v10.10, BD Biosciences). Data collection and analysis were not performed blind to the conditions of the experiments.

### AIM assay

Cryopreserved PBMCs were thawed in RF10, seeded at 1-2e6 cells per well of a 96-well plate, and rested for 4 hrs at 37°C. Alternatively, CCR6-depleted PBMCs via staining PBMCs with CCR6 PE-Cy7 (11A9, BD Biosciences) and anti-PE microbeads (Miltenyi Biotec) as per manufacturer’s protocol were seeded at 1-2e6 cells per well. Cells were stimulated either with 2 µg/ml of peptide (S_167_, S_751_), 1 µg/ml of peptide pool (S2), 1 µg/ml PepTivator SARS-CoV-2 Prot_S Complete peptide pool (Miltenyi Biotec), or an equivalent volume of DMSO (or sterile water as indicated) for 20 hrs. Selected experiments were also stimulated with SEB (5 µg/ml) as a positive control. CD154 APC-Cy7 (TRAP1, BD Biosciences) antibody was included in the culture medium for the duration of the stimulation. Cells were then washed in PBS and stained with Live/Dead Aqua and surface stained with the base T cell panel described above and the following antibodies: CD25 BB515 (M-A251, BD Biosciences), OX40 PerCP-Cy5.5 (Ber-ACT35, BioLegend), CD26 PE (BA5b, BioLegend), CD39 BUV395 (A1, BD Biosciences), CD30 PE (BERH8, BioLegend), CD30 PE (BY88, BioLegend), CD205 PE (DEC205, BioLegend), CD205 PE (HD30, BioLegend), CD205 APC (MMRI7, BD Biosciences), CXCR4 APC (12G5, BD Biosciences), CXCR4 PE (QA18A64, BioLegend). Optimal cell surface expression of CXCR4 required cells to be exposed to AIM assay culture conditions prior to antibody staining as previously reported ^30^. Cells were subsequently washed with 2% FCS/PBS, fixed with Cytofix (BD Biosciences), and data acquired with an LSR Fortessa (BD Biosciences).

### Transwell bystander activation cultures

PBMCs from the same individual were seeded at 2e6 cells in RF-10 in the upper and lower chambers of a 96-well Transwell unit separated by a 0.4 µm polycarbonate membrane pore Transwell insert (Corning). Anti-CD3/CD28 Dynabeads (Thermo Fisher) were added in the bottom chambers at a 5:1 PBMC:Dynabeads ratio. In some experiments, 2e6 T cells enriched from PBMCs via EasySep Human T Cell Isolation Kit (STEMCELL Technologies) were seeded in the top or bottom chambers of the Transwell culture. Where indicated, monoclonal antibodies for anti-human IL-2 (MQ1-17H12, BD Biosciences), anti-human IL-15 (34559, R&D Systems), and LEAF purified mouse IgG1κ isotype control antibody (BioLegend, MOPC-21) were added at 5 µg/ml in the bottom chambers. CD154 APC-Cy7 (TRAP1, BD Biosciences) antibody was included in the culture medium of both the bottom and top chambers of the transwell culture for the duration of the stimulation. After 20 hrs, the cells from each chamber were stained and analyzed separately according to the AIM assay. For permutations of new AIM combinations, where the average of the subtracted ‘AIM^+^’ values from the top chambers that were exposed to Dynabead-stimulated vs unstimulated PBMCs exceeded 0.2, these markers were considered prone to TCR-independent activation.

### Bulk RNAseq

PBMCs from the transwell bystander activation assay were sorted into the indicated populations by FACS into PBS. Cells were pelleted at 2000g for 10 mins, with supernatant aspirated to a final volume of 100 µL before resuspension and stored frozen at −80°C. Frozen cells were lysed directly in 3 volumes of QIAzol lysis reagent (QIAGEN). Genomic DNA removal and RNA extraction was performed using the Direct-zol RNA Microprep Kit (Zymo), according to the manufacturer’s protocol. The library preparation was performed by The Australian Genome Research Facility (Melbourne, Australia) using the Stranded Total RNA Prep with Ribo-Zero Plus kit (Illumina). Libraries were then sequenced using the NovaSeq 6000 for 150-bp paired reads. RNAseq analysis was performed using the web-based Galaxy platform maintained by Melbourne Bioinformatics ^59^. Reads were mapped to the *Homo sapiens* reference genome (hg38) using HISAT2 ^60^ and then quantified using HTSeq ^61^. Count matrices were generated and inputted into Degust (http://degust.erc.monash.edu) for differential gene expression using the limma (v3.40.6) method. The output was loaded into R for data visualization using *ggplot2* (v3.4.4). Genes were considered to be differentially regulated if the false discovery rate (FDR) was <0.05 and the absolute fold change (FC) was > 2.

### Single-cell RNA library preparation and sequencing

Cryopreserved PBMC from individuals receiving the IIV4 influenza vaccine (baseline, day 7 or day 28) were thawed and stimulated with recombinant HA proteins corresponding to the vaccine Ags. After 20 hrs of stimulation, cells were stained with a surface cocktail of antibodies (see above). Activated (OX40^+^) and resting (OX40^-^) memory (CD45RA^-^CXCR5^-^) and cTfh (CD45RA^-^CXCR5^+^) CD4^+^ T cells were sorted on a BD FACS Aria III. Cell hashing using anti-human hashtags C0251, C0253 and C0255 (BioLegend) were used to combine cells derived from different timepoints post vaccination into the same single-cell emulsion channel. Sorted cells were additionally stained with TotalSeq-C antibodies (BioLegend) against CD25 (BC96), OX40 (Ber-ACT35), and CCR6 (G034E3). A total of 30,000 cells were loaded onto a 10x Chromium controller and then prepared for sequencing using a Chromium Next GEM Single Cell 5’ Kit with feature barcoding and TCR immune receptor mapping (v2, Dual Index) from 10x Genomics. Libraries were generated according to the manufacturer’s protocol. The library quality was assessed with an Agilent 2100 Bioanalyzer and then sequenced on an Illumina NovaSeq 6000 platform with a depth of >20,000 reads per cell for gene expression libraries, and 5,000 reads per cell for VDJ and feature barcode libraries.

### Processing of single-cell RNA-seq data

Cell Ranger v6.0.1 ^62^ was used to process the gene expression and VDJ α/β FASTQ files with 10x Genomics refdata-gex-GRCh38-2020-A and refdata-cellranger-vdj-GRCh38-alts-ensembl-5.0.0 as references. Hashtag oligos (HTO) FASTQ files were processed with CITE-seq-Count v1.4.3. Output from CellRanger was analyzed in R using Seurat (v5.0.1) ^63^. Gene expression and hashtags were matched using the MULTIseqDemux ^64^, with cells with multiple hashtags excluded from downstream analysis. Data was subject to quality screening, gene expression was scaled and natural log-transformed, variable genes identified and dimensionality reduction using PCA for UMAP visualization. CD25^+^ and/or OX40^+^ clusters were demarked based on ADT-protein tags. TCR clonotype analysis was performed using *scRepertoire*. The complete workflow and associated scripts are available on [link available after acceptance].

### Statistical analysis

For all data other than RNA sequencing data, statistical analysis was performed in Prism 10 (GraphPad) using non-parametric statistical tests as indicated (making no assumptions about data normality). In all tests, statistical significance was quantified as \**p* < 0.5, \*\**p* < 0.01, \*\*\**p* < 0.001, and \*\*\*\**p* < 0.0001. No data were excluded from the study. Numerical values stated in the main text are given as the mean ± standard error of the mean (SEM).

## Acknowledgements

The authors thank the study participants for their involvement and provision of samples. We thank Kathleen Wragg, Grace Gare and Jane Batten for excellent technical assistance, and Danika Hill and Katherine Kedzierska for helpful discussion. We are grateful to Joanne Peterson, Joseph Sasadeusz and Helen Kent for the provision of samples from The Booster Study clinical trial. We thank Maggie Sakkas and Vanta Jameson at the Melbourne Cytometry Platform (Melbourne Brain Centre & Peter Doherty Institute node) for the provision of cell sorting services. We thank the NIH Tetramer Core Facility (contract number 75N93020D00005) for providing DPB1*04 monomers. This work was funded by an NHMRC Ideas grant to JAJ. HXT, SJK, AKW and JAJ are supported by NHMRC Investigator Grants, and JAJ is supported by the Sylvia and Charles Viertel Senior Medical Research Fellowship. The funders had no role in study design, data collection and analysis, decision to publish or preparation of the manuscript.

## Author contributions

Conceptualization: J.A.J

Methodology: M.Z.M.Z, H-X.T, A.H, A.K.W, J.A.J

Investigation: M.Z.M.Z, L.B, M-C.T, H-X.T, H.J.L, D.R., A.K.W, J.A.J

Visualization: M.Z.M.Z, L.B, H-X.T, A.K.W, J.A.J

Funding acquisition: H.X.T, S.J.K, A.K.W, J.A.J

Supervision: J.A.J, A.K.W, A.H

Writing-original draft: M.Z.M.Z, J.A.J

Writing-review & editing: M.Z.M.Z, H-X.T, S.J.K, J.A.J, A.K.W

All authors reviewed the final version of the manuscript.

## Data and materials availability

The sequencing datasets generated this study are available at GEO under access code [after acceptance].

---

**Figure S1.**
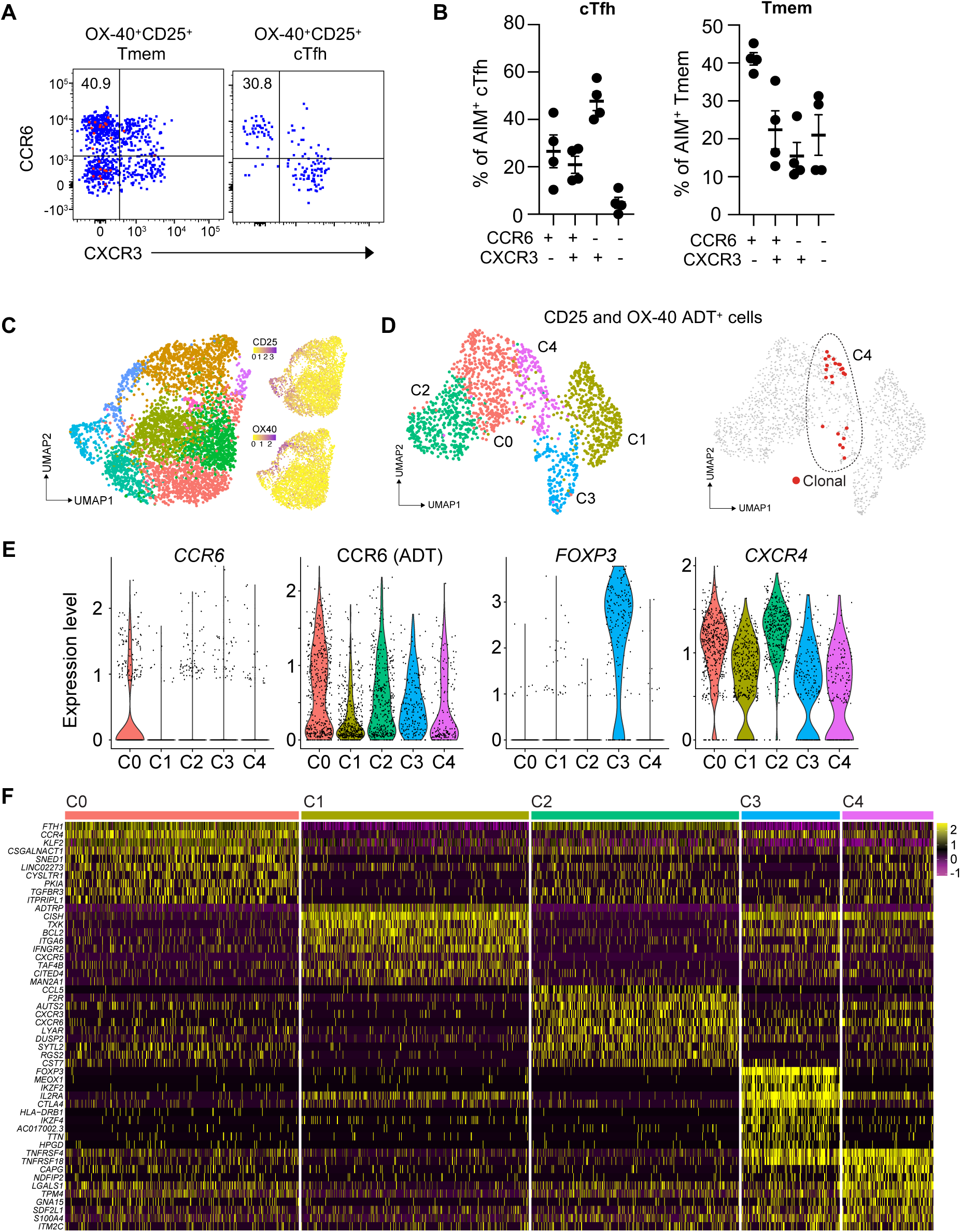
Transcriptional profile of human CD4^+^ Tmem cells after seasonal influenza immunization. **(A)** FACS plots and **(B)** frequencies of CCR6/CXCR3 expression on AIM^+^ CD4^+^ Tmem or cTfh following *in vitro* HA protein stimulation. PBMC samples were collected 7d post-immunization (n=4 individuals). **(C)** UMAP plots of CD4^+^ T cells sorted from PBMCs (n=3 participants) at d0, 7, 21 post-immunization alongside ADT-defined surface expression of CD25 and OX40. **(D)** UMAP plots of activated (CD25^+^ and/or OX40^+^ ADT expression) memory CD4^+^ T cells with positioning of expanded TCR CDR3 clonotypes shared between multiple cells. **(E)** Expression of indicated genes in unbiased clusters. **(F)** Heatmap depicting the top DEGs identified within each cluster.

**Figure S2.**
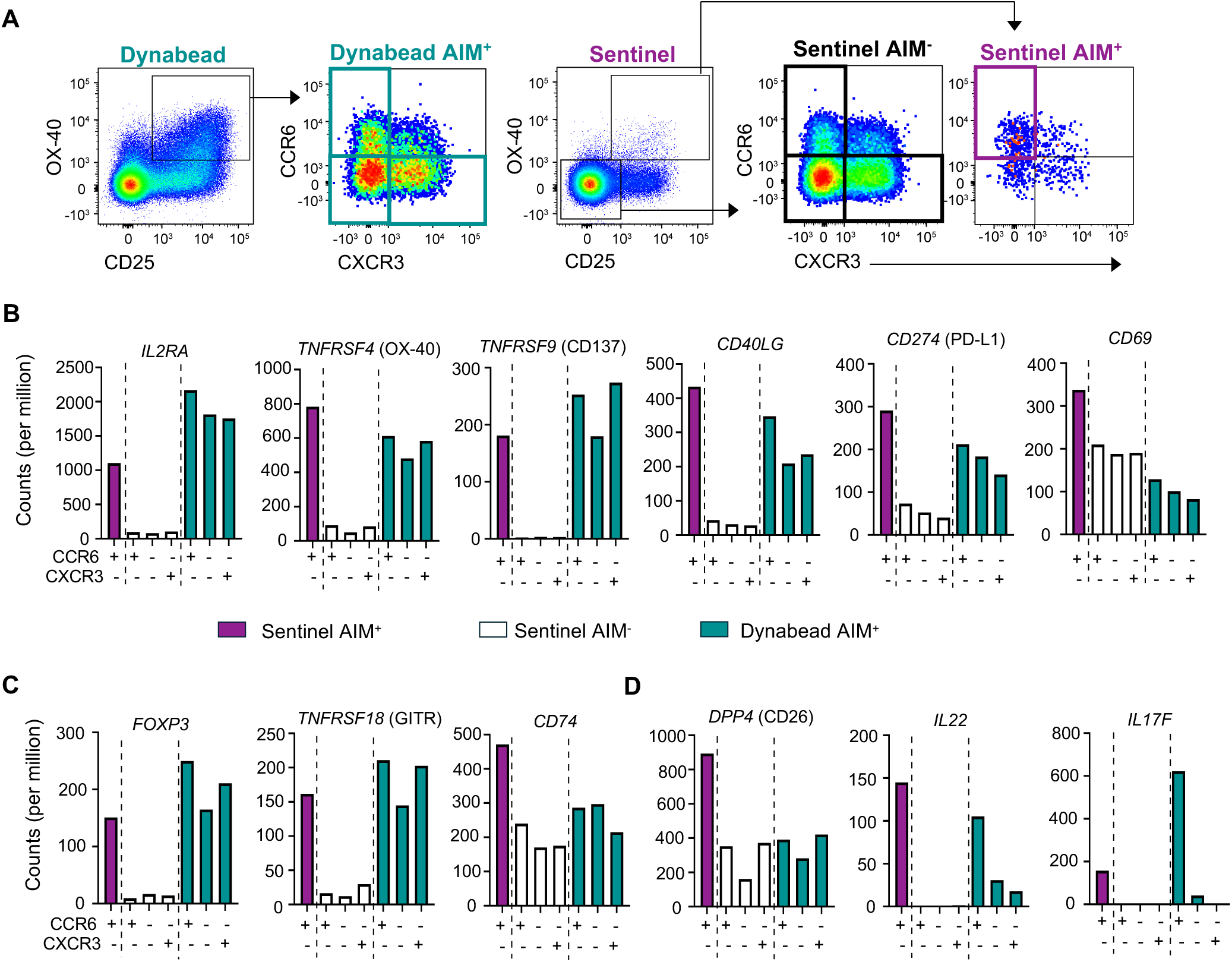
Differential gene expression analysis of sentinel AIM^+^ CCR6^+^ T cells in the transwell model. **(A)** FACS plots depicting sorted populations of AIM^+^ or AIM^-^ cells from Dynabead and sentinel wells for bulk RNAseq. **(B)** Gene expression for the commonly used AIM markers *IL2RA* (CD25), *TNFRSF4* (OX40), *TNFRSF9* (CD137/4-1BB), *CD40LG* (CD154), *CD274* (PD-L1), and *CD69* (CD69). **(C)** Expression of the Treg-associated genes *FOXP3* (Foxp3), *TNFRSF18* (GITR), and *CD74* (CD74). **(D)** Expression of the Th17/Th22-related genes *DPP4* (CD26), *IL22* (IL-22), and *IL17F* (IL-17F).

**Figure S3.**
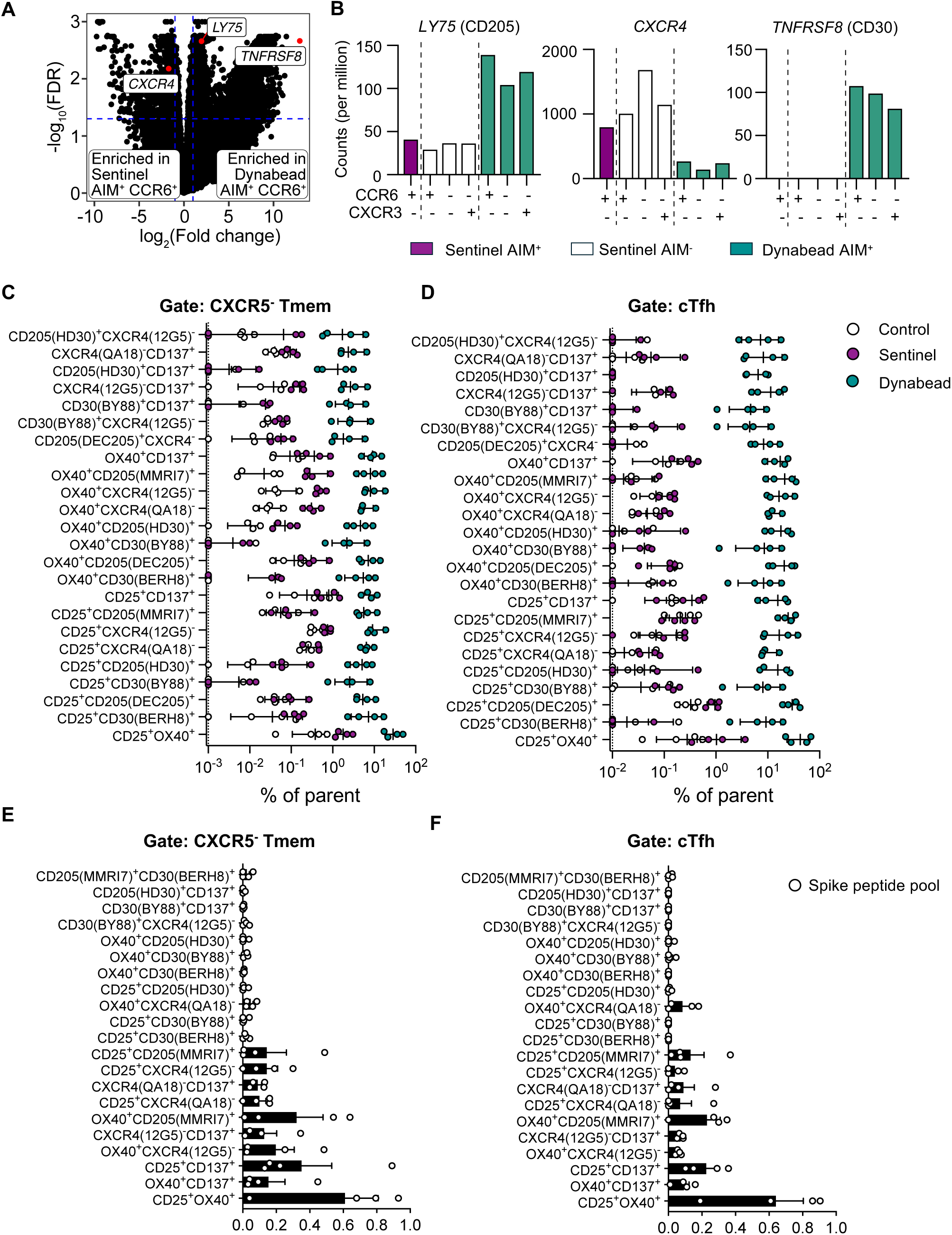
Bulk RNAseq identifies new AIM candidates that distinguish T cells activated in a TCR-independent or -dependent manner. **(A)** Volcano plot of selected genes enriched in Dynabead AIM^+^ CCR6^+^ relative to sentinel AIM^+^ CCR6^+^ CD4^+^ T cells. Dotted lines indicate criteria threshold of -log_10_(0.05) FDR and at least a log_2_ fold change of −1 or 1. n=1 individual. **(B)** Expression of *LY75* (DEC-205/CD205), *TNFRSF8* (CD30), and *CXCR4* (CXCR4/CD184). **(C-D)** Frequencies of AIM^+^ cells for marker combinations with low or no TCR-independent activation (see Materials & Methods for low threshold criteria) for (**C)** CD4^+^ Tmem or **(D)** cTfh cells in the transwell assay (n=4 individuals). **(E-F)** Frequencies of AIM^+^ (DMSO subtracted) cells among **(E)** CD4^+^ Tmem and **(F)** cTfh cells following *in vitro* stimulation of SARS-CoV-2 vaccinated/convalescent PBMCs (n=4) with an S2 peptide pool.

